## Supplementary file_5 for "DANCE: An open-source analysis pipeline and low-cost hardware to quantify aggression and courtship in *Drosophila*"

**Supplementary File 5: Quantifying identity switch rates across recording setups**

| **Video Set** | **No. of videos** | **Frame rate** | **Total frames per video** | **No. of identity switches** | **Frames with identity switch** | **Total frames** | **Error rate** |
| --- | --- | --- | --- | --- | --- | --- | --- |
| High resolution (with decapitated virgin videos) | 19 | 30 | 27068 | 7 | 3425 | 5142912 | 0.66402 |
| ‘DANCE’ Medicinal blister packs (with decapitated virgin videos) | 24 | 30 | 27068 | 20 | 12354 | 64962 | 1.901692 |

We used the **Visualizer** tool within FlyTracker to flag potential identity swaps, identified when flies overlap (occlusions), make sudden jumps, or move unusually close to one another. **These criteria were intentionally broad to capture all possible identity errors.** This approach minimizes manual inspection by restricting it to flagged intervals rather than reviewing every frame.

We focused on **decapitated virgin videos**, as this condition poses the greatest challenge for maintaining fly identity: males and decapitated females are similar in size, increasing the likelihood of swaps. Using this approach, we compared two datasets. **High-resolution machine-vision recordings** showed **7 identity switches across 19 videos (error rate = 0.66%)**, whereas **DANCE recordings using standard smartphone cameras** had **20 switches across 24 videos (error rate = 1.9%).**

Although the phone-camera recordings produced slightly more identity switches than the high-resolution system, the modest increase in error demonstrates that **accurate and reliable tracking can be achieved even with low-cost, accessible hardware.**
